# Effects of species traits and ecosystem characteristics on species detection by eDNA metabarcoding in lake fish communities

**DOI:** 10.1101/2020.09.25.314336

**Authors:** Hideyuki Doi, Shunsuke Matsuoka, Shin-ichiro S. Matsuzaki, Mariko Nagano, Hirotoshi Sato, Hiroki Yamanaka, Saeko Matsuhashi, Satoshi Yamamoto, Toshifumi Minamoto, Hitoshi Araki, Kousuke Ikeda, Atsuko Kato, Kouichi Kumei, Nobutaka Maki, Takashi Mitsuzuka, Teruhiko Takahara, Kimihito Toki, Natsuki Ueda, Takeshi Watanabe, Kanji Yamazoe, Masaki Miya

**Author notes:** These authors contributed equally to this study (Hideyuki Doi, Shunsuke Matsuoka, Shin-ichiro S. Matsuzaki). Corresponding author: Hideyuki Doi.

## Abstract

Although environmental DNA (eDNA) metabarcoding is acknowledged to be an exceptionally useful and powerful tool for monitoring surveys, it has limited applicability, particularly for nationwide surveys. To evaluate the performance of eDNA metabarcoding in broad-scale monitoring, we examined the effects of species ecological/biological traits and ecosystem characteristics on species detection rates and the consequences for community analysis. We conducted eDNA metabarcoding on fish communities in 18 Japanese lakes on a country-wide scale. By comparing species records, we found that certain species traits, including body size, body shape, saltwater tolerance, and habitat preferences, influenced eDNA detection. We also found that the proportion of species detected decreased significantly with an increase in lake surface area, owing to an ecosystem-size effect on species detection. We conclude that species traits, including habitat preferences and body size, and ecosystem size should be taken into consideration when assessing the performance of eDNA metabarcoding in broad-scale monitoring.

## Introduction

An ongoing global decline in biodiversity is beyond doubt and is particularly pronounced in freshwater ecosystems^1–3^. Nevertheless, despite concerns regarding the loss of local/global freshwater fish diversity^1,2,4^, there has been a recent tendency to scale back on investments in monitoring and museum collections^5,6^. An understanding of the prevailing status and trends of biodiversity provides a bedrock for ecological research and conservation biology^1,2^, and in this regard, Matsuzaki et al.^7^ highlighted that long-term trends in local-scale fish diversity have not been comprehensively monitored at broader scales (i.e., national or continental scales) and that performing broad-scale surveys of fish communities is essential for planning fish management and conservation measures^8^. However, the number of lake fish surveys conducted by national and local governments have recently declined^5,6,9^. Given this undesirable trend, it would be prudent to develop broad-scale survey methods for fish monitoring, which could be employed in conducting national-level surveys and thereby provide a basis for formulating conservation policies.

Recent advances in molecular ecology have seen the emergence of environmental DNA (eDNA) analysis as a useful approach for investigating the distribution and richness of aquatic and terrestrial organisms^10–16^, and high-throughput parallel DNA sequencing has recently been applied to eDNA methods for simultaneous detection of multiple taxa, known as eDNA metabarcoding^16–21^. For example, Miya et al.^22^ designed and applied universal PCR primers (MiFish primers) for fish eDNA metabarcoding, and MiFish primers recently developed for different fish taxa^15,16,23^ have shown higher performance compared with other primers types^24^ and PCR conditions^25^.

eDNA metabarcoding is acknowledged to be an exceptionally useful and powerful tool for community surveys^16–20,26,27^, and consequently, in recent years, this technique has been widely applied in aquatic community surveys worldwide^28^. However, despite the growing number of eDNA metabarcoding studies being conducted, the performance of eDNA metabarcoding applied in broad-scale (e.g., national-wide) surveys has, with a few exceptions^18,26,29^, yet to be sufficiently evaluated quantitatively and statistically^28^.

From the perspective of evaluating the performance of eDNA metabarcoding surveys, we hypothesized that the ecological/biological traits of targeted species and ecosystem characteristics would influence the detection of species when using eDNA metabarcoding; based on the fact that fish ecological/biological traits (e.g., body size and habitat preference) have been considered to be factors that affect the detectability of eDNA^30,31^. In this regard, Pont et al.^31^ revealed differences in the traits of fish detected in a large river using eDNA metabarcoding and traditional survey methods. Differences in the relative abundance of fish related to habitat preference (benthic or pelagic), dietary type, and level of tolerance to water pollution have also been detected in comparisons between eDNA metabarcoding and traditional surveys^31^. The findings of these studies indicate that the ecological/biological traits of fish species, such as body size and shape, habitat preference, and environmental tolerance, may influence specific detection rates when using eDNA metabarcoding. However, despite the increasing evidence of the effects of species traits on the detectability of eDNA, assessments of these effects across spatially broad areas, for example, entire countries with diverse species, are still minimal. Given the same sampling effort, the detection of species eDNA may also depend on certain ecosystem characteristics, such as ecosystem size (area and volume) and habitat types and variation. For example, eDNA metabarcoding may achieve a lower rate of species detection in larger lakes where greater sampling effort is required to characterize fish communities.

In an effort to gain a better understanding of the factors that could potentially influence to the efficacy of eDNA metabarcoding, we conducted surveys of fish communities in 18 lakes distributed throughout Japan using universal MiFish primers^22^ and compared the results obtained with the records of over 200 fish species (including non-native species) inhabiting these lakes^7^. Based on this comparison, we were able to assess the validity of the aforementioned hypotheses regarding the use of eDNA metabarcoding surveys. Moreover, to evaluate the utility of different sampling methods for broad-scale eDNA surveys, we also conducted a survey using individual and mixed water sampling techniques and different cooling and freezing methods to evaluate the effects of sampling and transportation on the findings of eDNA surveys. Finally, we discuss the applicability of eDNA metabarcoding for broad-scale monitoring in lake fish, taking into consideration species traits and ecosystem characteristics.

## Results

### Sequencing and eDNA metabarcoding overview

Given that we obtained a sufficient proportion of sequences from the raw data and that, with the exception of Lake Shikotsu, there were very few non-fish sequences (Table S2), we assumed MiSeq sequencing and pipeline analysis of the sequence data to have been successful. MiSeq paired-end sequencing for library construction (N = 178: 144 samples, 18 field and filter blanks, and 16 PCR negative controls) yielded a total of 5,208,062 sequences [29,258 ± 23,237 (mean ± SD) sequences for each sample, Table S2].

### Number of detected fish taxa

In total, 119 fish taxa were detected based on eDNA metabarcoding, and all detected taxa were identified to the species or genus level (Table S4, Figs. 1 and S3). We found significant differences among the different sampling methods with respect to the number of fish taxa detected using eDNA metabarcoding (nested ANOVA with LMM, P < 0.001, Table S5 and Fig. S4). However, we also detected six species in 23 of the negative and field controls. We obtained sequence reads from six species (*Carassius auratus, Chelon haematocheilus, Micropterus salmoides, Pagrus major, Rhinogobius* spp., and *Tribolodon hakonensis*, Table S4). Furthermore, five species found in the fish records had lower sequence reads (sequences < 276) in the controls than in the samples. In contrast, *Pagrus major* (Red seabream) was not found in the records and does not inhabit the surveyed lakes. The number of sequence reads obtained for *P. major* (sequences = 29 and 452) were found to be higher than those species in the samples (sequences= 20 and 21). Thus, we assumed that the species DNA was contaminated and excluded the species from further analyses.

**Figure 1.**
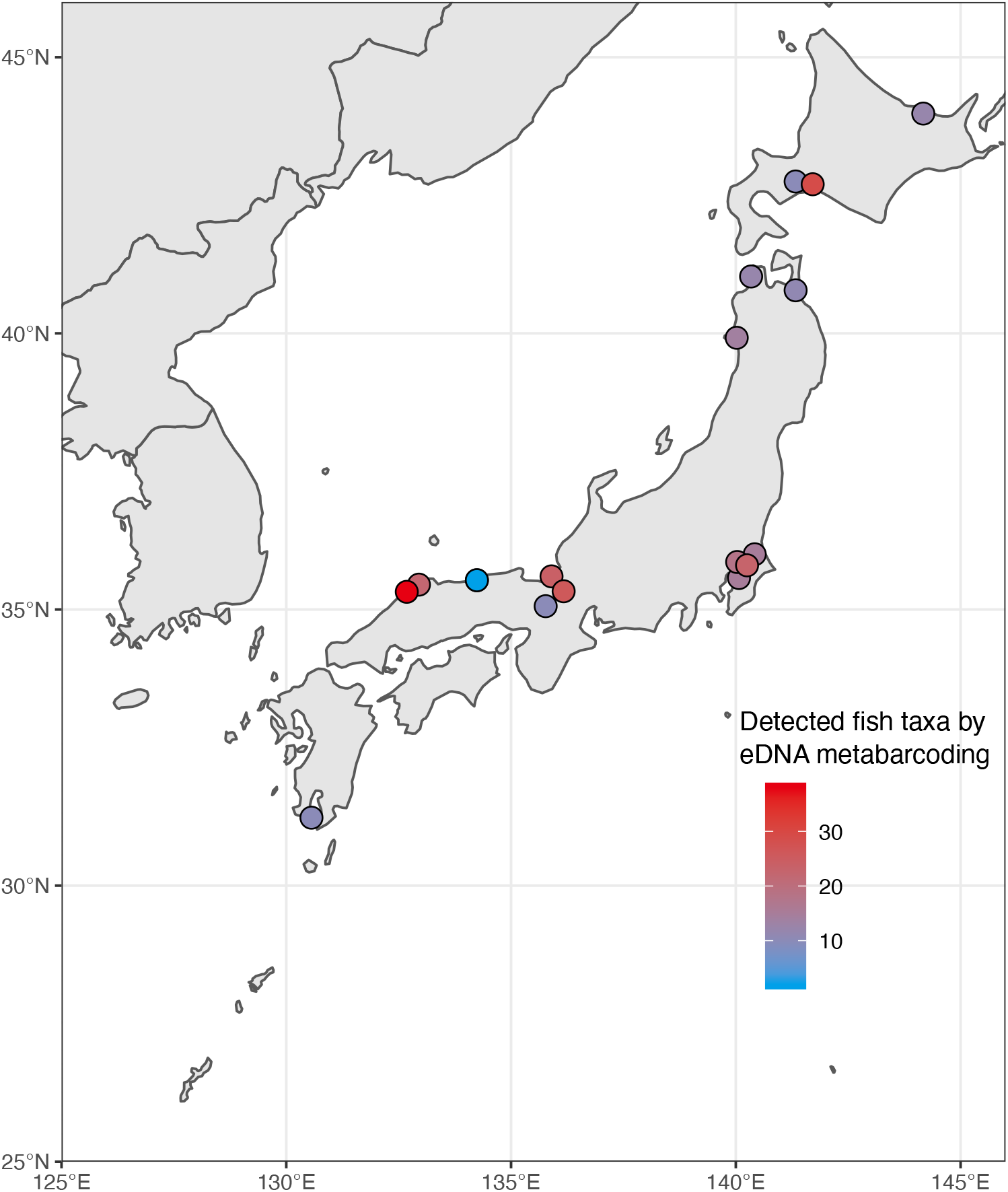
A map of Japan showing the locations of the 18 study lakes. The color of circles indicates the approximate numbers of fish taxa detected using eDNA metabarcoding based on the Mix_cool sampling method. Total numbers of fish taxa detected using other sampling methods (Individual and Mix_freeze) are shown in Fig. S3.

We found differences in the number of fish taxa in the different lakes (Figs. 1 and S5), and the number of fish taxa detected using the Mix_cool and Mix_freeze methods was significantly higher than those detected based on individual sampling (Fig. 2a, nested ANOVA with LMM, P < 0.001 and Tukey’s multiple comparison, Table S5). Nevertheless, we found that some species detected in the individual samples were not detected in the Mix_cool and Mix_freeze samples (Table S4).

**Figure 2.**
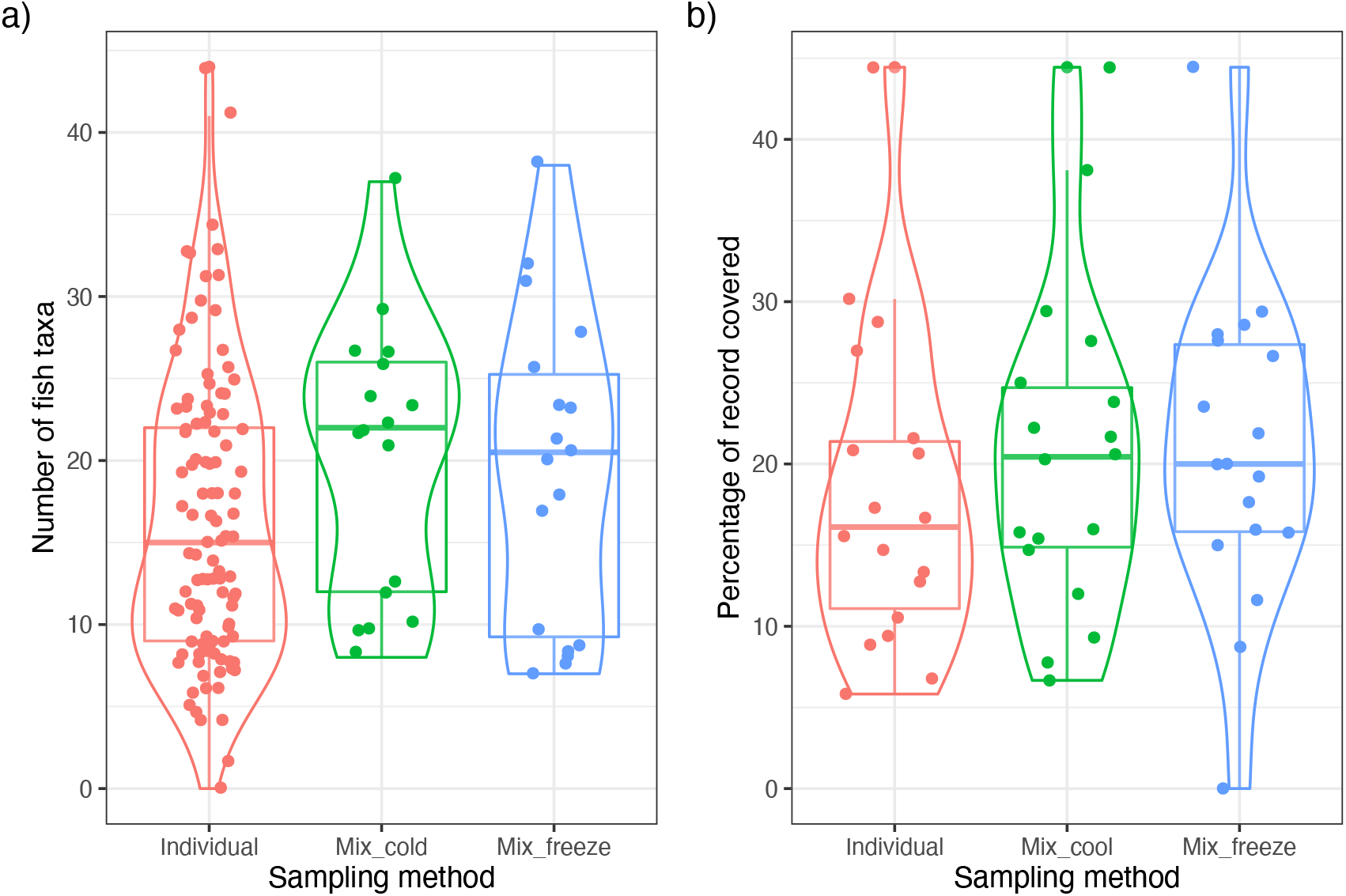
Box and violin plots showing (a) the fish taxa detected and (b) the percentage of record covered (%) with eDNA metabarcoding using different sampling methods. The boxes and bars in the box plots indicate median ± inter-quartiles and ±1.5 × inter-quartiles, respectively. The points represent individual data values. The smooth lines indicate the distribution of the data using violin plots. The violin plot outlines illustrate kernel probability density, i.e., the width of the enclosed area represents the proportion of the data located.

### Percentage record covered

To evaluate the species detection performance of eDNA metabarcoding, we used “percentage record covered (%)” as a measure to indicate the proportion of eDNA-detected species (number of species) matching those in the species records. We accordingly found that the percentage record covered did not differ significantly among the three sampling methods (Fig. 2b, nested ANOVA with LMM, P = 0.249, Table S5), although values were found to differ among the surveyed lakes (Fig. S6).

We also compared the ecological/biological traits of species groups frequently detected using eDNA metabarcoding or in the species records, which was evaluated based indicator taxa analysis (Fig. 3). All the nested ANOVAs with linear mixed models (LMMs) for the fish traits revealed significant differences between eDNA metabarcoding and the species records, whereas no significant differences were detected among the different sampling methods (Table S6). We found that species detected at a higher frequency using eDNA metabarcoding tended to have shorter body lengths (Fig. 3a), mainly inhabit the benthopelagic zone (Fig. 3b), and are saltwater tolerant (Fig. 3c). With respect to lateral body shape types, smaller proportions of eel-like species, either short, deep bodied, or both, were identified among those species detected with a higher frequency using eDNA metabarcoding than in the species records (Fig. 3d).

**Figure 3.**
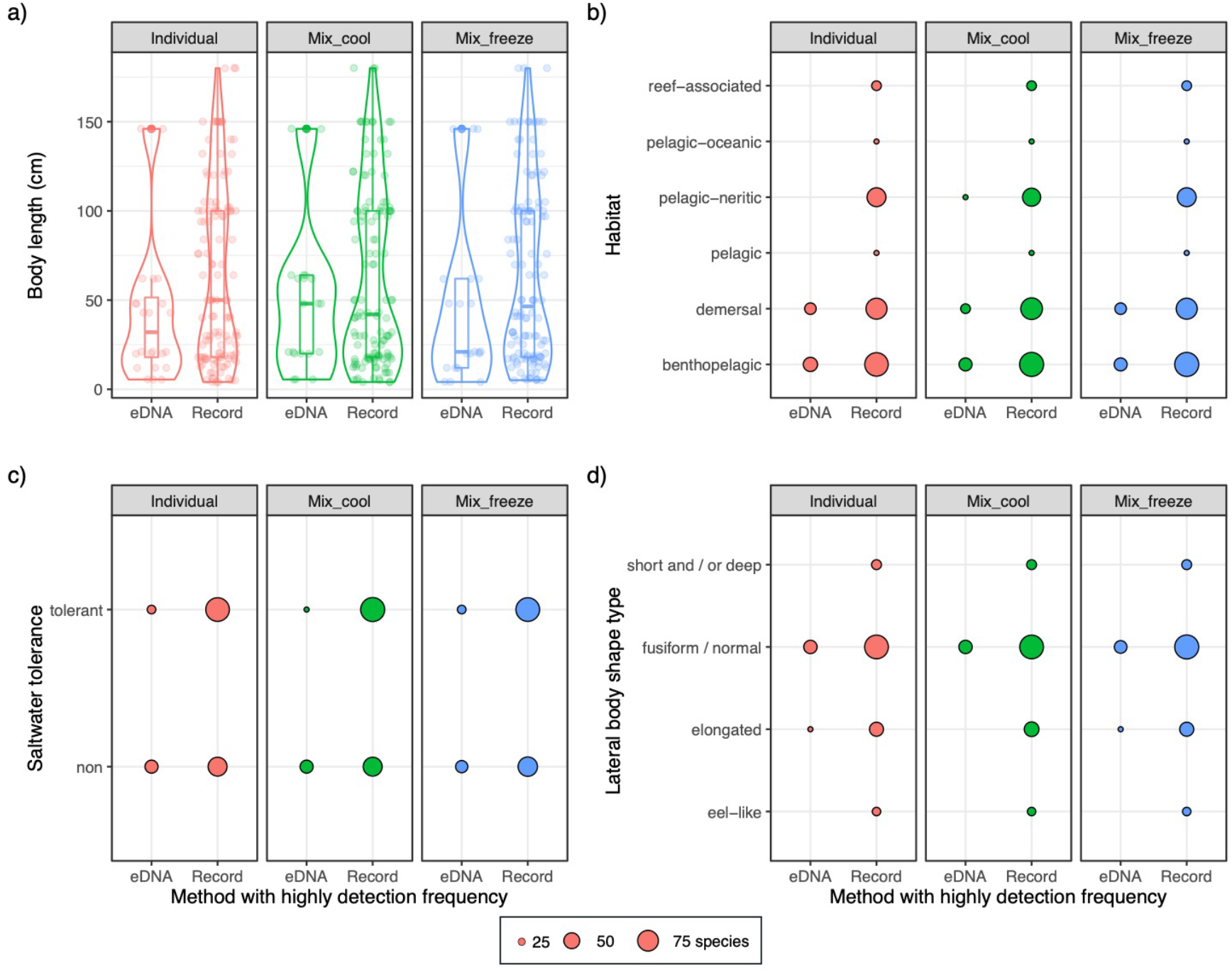
The ecological/biological traits of fish species in species group most frequently detected based on eDNA metabarcoding and in the species records using the different sampling methods, as evaluated based on indicator taxa analysis. (a) Box and violin plots for body length. The boxes and bars in the box plots indicate median ± inter-quartiles and ±1.5 × inter-quartiles, respectively. The smooth lines indicate the distribution of the data using violin plots. The violin plot outlines illustrate kernel probability density, i.e., the width of the enclosed area represents the proportion of the data located. The points represent individual data values. (b-d) Balloon plots indicating the habitat preference (b), saltwater tolerance (c), and lateral body shape type (d) of fish species. The sizes of balloons in (b), (c), and (d) indicate the number of species in each category, as shown in the bottom legend.

To evaluate the effect of lake limnological features on the percentage record covered, we used generalized linear models (GLMs), which revealed a significant negative effect of surface area on the percentage record covered (P < 0.018, Table S7, Fig. 4), whereas for all sampling methods used, there were no significant differences with respect to latitude, mean water depth, trophic state, or water type (P > 0.05, Table S7). The relationship between the percentage record covered and the non-included factors to the final GLMs are shown in Fig. S7.

**Figure 4.**
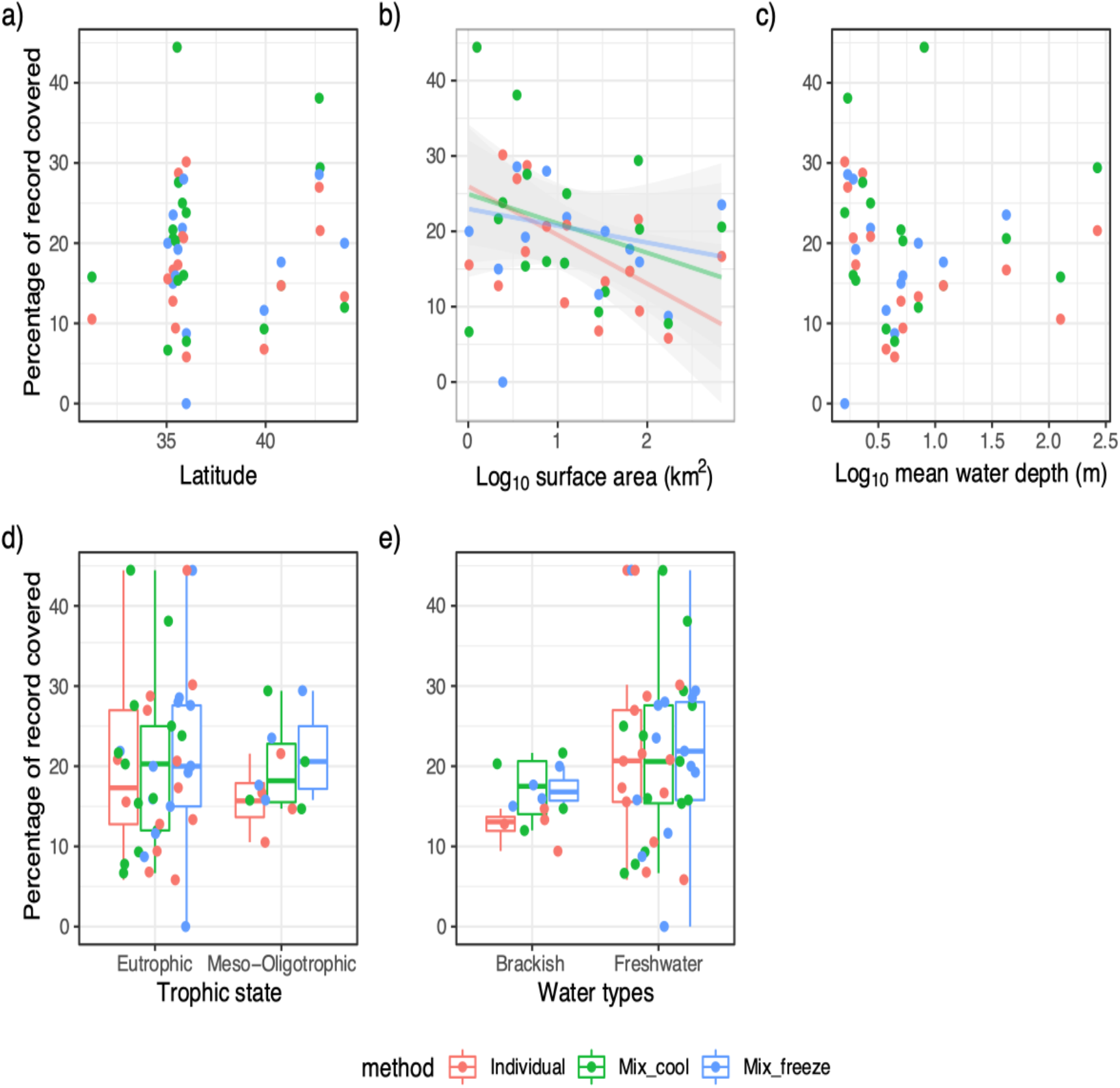
The relationships between the percentage of record covered by eDNA metabarcoding (%) against the fish records and (a) lake latitude, (b) lake surface area, (c) lake mean depth, (d) trophic state, and (e) water type, including the final GLMs, using the different sampling methods (red = Individual, green = Mix_cool, and blue = Mix_freeze). Plots show the relationships with other factors are shown in Fig. S8. The solid and gray areas indicate the regression line from the significant GLM results and the 95% CI, respectively. The boxes and bars in the box plot indicate median ± inter-quartiles and ±1.5 × inter-quartiles, respectively. The points represent individual data values.

### Community analysis

Using indicator taxa analysis, we established that certain species were detected at a significantly higher frequency using eDNA metabarcoding than indicated in the species records for the 18 surveyed lakes (please refer to Table 1 for species with significantly higher frequencies determined by eDNA metabarcoding and all results in Table S8). For example, *C. auratus* was significantly more frequently detected using all three sampling methods, whereas *Zacco platypus* was significantly more frequently detected in the individual and Mix_freeze samples, *Hemibarbus labeo* and *Hypophthalmichthys nobilis* were more frequently detected in individual samples, and *Rhinogobius* spp. were significantly more frequently detected in Mix_freeze samples.

**Table 1.** Significantly higher frequencies of species detected using eDNA metabarcoding compared with species records, as determined based on indicator taxa analysis (P < 0.05). “Best” indicates the method facilitating the most frequent detection. P values were calculated based on 999 permutations subsequent to Sidak’s correction for multiple testing. Species detected at non-significant, and significantly lower frequencies in the species records are shown in Table S7.

| Method | Species name | P value for eDNA | P value for species records | Best method | P value for multiple testing |
| --- | --- | --- | --- | --- | --- |
| Individual | *Carassius auratus* | 0.001 | 1 | eDNA | 0.001999 |
| Individual | *Hemibarbus labeo* | 0.001 | 1 | eDNA | 0.001999 |
| Individual | *Hypophthalmichthys nobilis* | 0.001 | 1 | eDNA | 0.001999 |
| Individual | *Zacco platypus* | 0.001 | 1 | eDNA | 0.001999 |
| Mix_cool | *Carassius auratus* | 0.001 | 1 | eDNA | 0.001999 |
| Mix_cool | *Zacco platypus* | 0.023 | 0.997 | eDNA | 0.045471 |
| Mix_freeze | *Carassius auratus* | 0.001 | 1 | eDNA | 0.001999 |
| Mix_freeze | *Rhinogobius* spp. | 0.023 | 0.997 | eDNA | 0.045471 |

Non-metric multidimensional scaling (NMDS) ordination analysis revealed dissimilarities in fish communities among the surveyed lakes and between eDNA metabarcoding and the species records (Fig. 5 for Mix_cool and Figs. S8 and S9 for individual and Mix_freeze, respectively). Similarly, PERMANOVA analysis revealed significant differences among the lakes and between eDNA metabarcoding and the species records (P < 0.003, Table S9).

**Figure 5.**
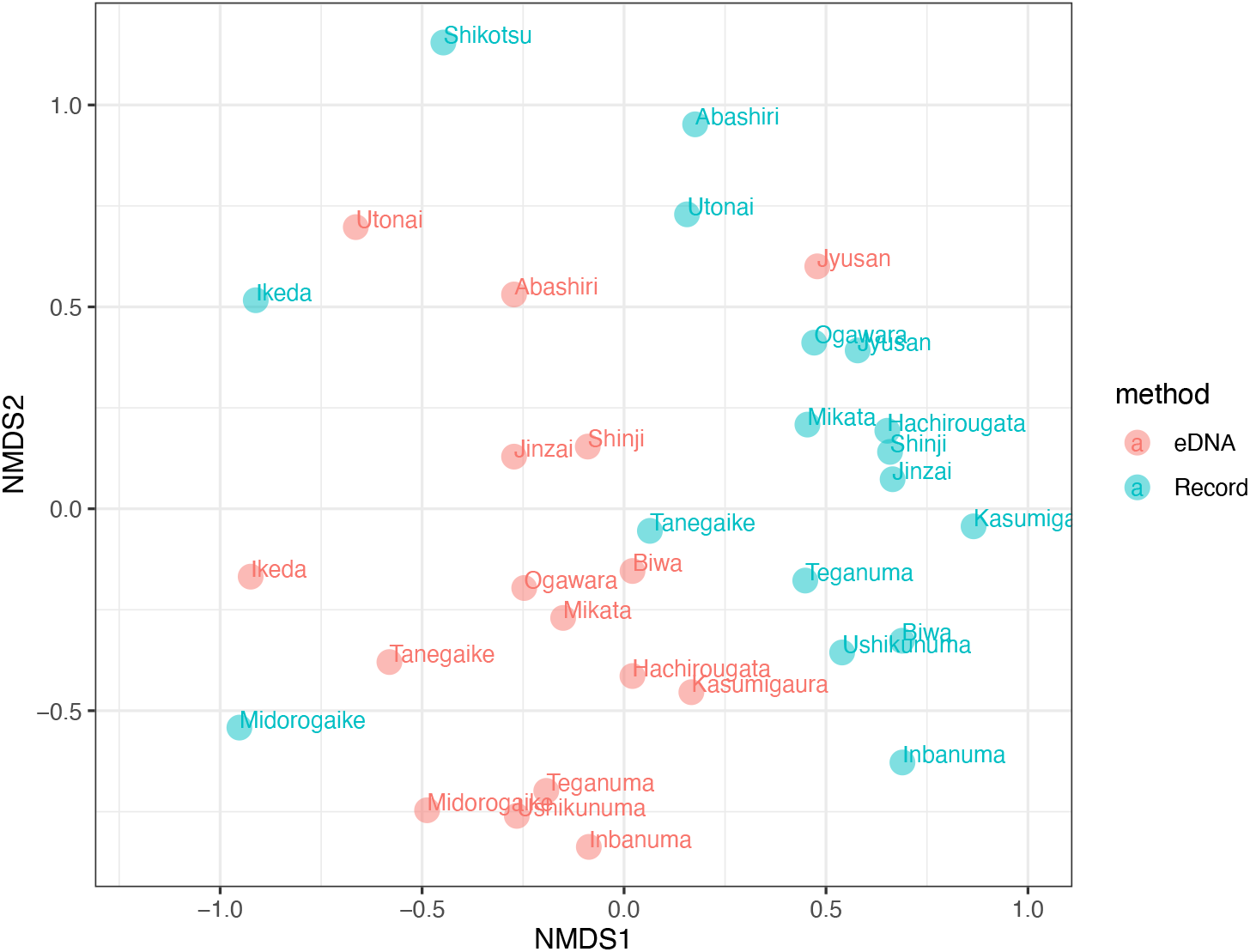
A non-metric multidimensional scaling (NMDS) ordination plot for fish communities based on species records and eDNA metabarcoding using Mix_cool samples. The NMDS stress was 0.174. NDMS plots for Individual and Mix_freeze data are shown in Figs. S8 and S9, respectively.

## Discussion

In this study, we observed a difference in the ecological/biological traits of fish species and lake limnological features relating to the eDNA-evaluated community structure and species detection based on broad-scale eDNA and direct surveys.

We initially compared the efficacy of three different sampling methods (individual, Mix_cool, and Mix_freeze sampling) for water collection, and accordingly found that the number of fish taxa detected using Mix_cool and Mix_freeze methods was significantly higher than that detected using individual sampling. In contrast, we detected no significant differences among these sampling methods based on the percentage of record covered, NMDS, or indicator taxa analyses. In this regard, Sato et al.^20^ suggested that mixed samples can be used for fish community comparisons despite a somewhat lower detection rate for certain rare species, and this indeed appeared to hold true with respect to the lakes we sampled in the present study.

Our findings tended to indicate that the water transportation method (cool vs. frozen) had no significant influence on the efficiency of eDNA detection, which is consistent with the findings of Deiner et al.^32^, who reported a non-significant difference between cool and frozen water transportation on eDNA detection by quantitative PCR (qPCR), which is in contrast to the findings of Takahara et al.^33^, who found a reduction in the fish species detected using frozen preservation of water samples based on qPCR analyses. We thus believe that both methods may be useful for transporting water samples for eDNA analysis. In this regard, as a consequence of a field survey we conducted in 2015, Yamanaka et al.^15^ developed a water preservation method for eDNA analysis that included the addition of the quaternary ammonium compound benzalkonium chloride (BAC), and we recommend using this BAC method for water transportation.

In order to evaluate the performance of eDNA metabarcoding, we examined the percentage record covered for all three sampling methods. We accordingly found that the values obtained for species records were approximately 30%, thereby indicating that many of the species inhabiting the surveyed lakes, particularly pelagic species and those with lower occurrence, would not be detected using eDNA metabarcoding. Below, we consider the factors (limnological features and species traits) that might contribute to the lower detection rates obtained using eDNA metabarcoding.

As factors that could potentially influence the efficacy of eDNA metabarcoding, we examined the selected ecological/biological traits of fish species, and accordingly found that species with shorter body length and normal body shape, those inhabiting benthopelagic habitats, and those showing tolerance to saltwater were frequently detected using eDNA metabarcoding. In previous eDNA studies, certain biological traits of fish, notably body size, have been considered to influence eDNA detection^30,31^. However, whereas the findings of several experimental studies have indicated that fish of larger body size release larger quantities of eDNA^30^, we found that species with shorter body length, as an index of body size, were detected more frequently using eDNA metabarcoding than their occurrence in the species records. We suspect that these differences are probably attributable to the size–abundance distribution of fish communities, which indicates that species with smaller body sizes are more abundant in lakes than are those of larger body size^34^. Lakes, particularly those characterized by brackish waters, harbor numerous marine-migratory fish species that show tolerance to saltwater, and eDNA metabarcoding surveys of lakes have detected many such migratory species^16,27^. Given that the timing of occurrence of species influences the detection of their eDNA^35^, the timing of migration would influence the detection of migratory species in lakes using eDNA metabarcoding, and consequently, may contribute to the infrequent detection of saltwater-tolerant species by eDNA metabarcoding in eDNA sampling surveys. Spatial heterogeneity in fish eDNA concentrations/detection in aquatic ecosystems has previously been observed^10,15,25,26^, and Pont et al.^31^ found that benthic species were detected at a higher rate using eDNA metabarcoding than by using traditional methods. This phenomenon would appear to suggest that the habitat preferences of fish might influence their detectability using eDNA metabarcoding. In this regard, the lakeshore water sampling conducted in the present study revealed the presence of numerous benthopelagic species based on eDNA metabarcoding, but relatively few pelagic species. Accordingly, in order to detect a larger number of pelagic species, we would need to perform offshore sampling, for example, using drone-assisted water collection for eDNA surveys^36^. The lateral body shape traits of species may also be associated with their micro-habitat use; for example, species with eel-like and short and/or deep body shapes appear to prefer sheltered micro-habitats (i.e., beneath stones) and pelagic zones. Consequently, the preference for macrohabitats, such as the benthic and pelagic zones, and their microhabitat use with respect to body shape types could serve as useful indices for evaluating eDNA detectability.

Evaluation of the effects of lake characteristics on the percentage record covered based on our modeling results revealed significant negative effects of lake surface area. Lakes with a larger ecosystem size are characterized by a broad diversity of habitats types, and correspondingly tend to support a higher diversity of fish species^37,38^ and an increased proportion of pelagic zones^39^. Consequently, the size of lake ecosystems will tend to influence the rate of detection using eDNA metabarcoding. With respect to rivers, Bylemans et al.^17^ found that river morphology influences the optimal sampling strategy for eDNA metabarcoding in these habitats, and in the present study, we identified a similar effect of ecosystem morphology on eDNA sampling strategy in lakes. Taking into consideration the factors of sampling effort and expenditure, broad-scale surveys, such as nationwide surveys, tend to be constrained with respect to limited sample sizes, and given these circumstances, it may thus be important to assess the performance of eDNA metabarcoding-based limited sampling effort. In this regard, our GLM analysis predicted that the percentage record covered would decline to almost half (for example, from 30% to 15%) with an increase in lake surface area from 0.1 to 100 km^2^.

The utility of eDNA metabarcoding in evaluating fish community structure in lakes has been examined in previous studies that have compared eDNA metabarcoding and traditional sampling methods^16,26^. Similar to the previous results, we observed significant differences in the fish communities detected using eDNA metabarcoding and those in the species records. As discussed above, lake ecosystem size and the traits of fish species can influence the results obtained using eDNA metabarcoding, and consequently, we would expect these effects to be reflected in differences between fish community structures determined based on eDNA metabarcoding and the species record. We indeed found that similar to that shown in the species records, the fish community structure determined by eDNA metabarcoding showed significant differences among the surveyed lakes, thereby indicating that community analysis using this technique could compare the communities of different lakes. Although eDNA metabarcoding would be useful for broad-scale community surveys at low survey costs, we should carefully consider the influence of ecosystem characteristics and the traits of species within fish communities in such surveys.

We acknowledge that the present study does have certain limitations, and thus the conclusions we draw should be viewed as provisional and in need of further verification. Notably, as we performed only single samplings for the eDNA survey, we were unable to evaluate the false negative/positive detection rates for fish species, which is a vital consideration in applying eDNA metabarcoding to biomonitoring^40^. eDNA sampling strategies are also prone to false-positive errors, owing to contamination and/or errors in PCR or sequencing, which may result in the spurious detection of species^41,42^. Accordingly, further studies are needed to confirm false negative/positive detection rates in lake eDNA surveys.

In conclusion, on the basis of a comparison with existing fish records, we were able to establish that certain traits of fish species and characteristics of ecosystems, particularly body size/shape, species habitat preferences, and ecosystem size, would be useful indices for optimizing eDNA metabarcoding sampling strategies, as well as for assessing the rates of species detection using eDNA methods. From the perspective of designing eDNA metabarcoding-based surveys to monitor aquatic communities across wide geographical areas, we should consider the ecological/biological traits of species and the characteristics of an ecosystem that can potentially influence species detectability.

## Methods

### Study lakes and sampling sites

For the survey described herein, we selected 18 lakes distributed throughout Japan (Fig. 1), the locations of which are listed in Table S1. The lakes differed with respect to the surface area, water depth, volume, trophic state, and water type (brackish and freshwater), as shown in Table S1. Along the shores of each lake, we established six sampling sites separated by approximately equal distances (Fig. S1), none of which were located in the vicinity of river inflows/outflows.

As species-area accumulation curves in community surveys^43,44^, increasing the sample size would increase the detectability of species by eDNA metabarcoding^20,45^. Given that one of our aims in this study was to evaluate the effect of lake size on the detectability of species using eDNA metabarcoding, we conducted surveys with an equal number of sampling sites regardless of lake size.

### Collection water samples for eDNA survey

Between 14 July and 4 November 2015, we collected samples of water at each of the six sampling sites located along the shores of the surveyed lakes, using three sampling methods, namely, “Individual,” “Mix_cool,” and “Mix_freeze” (Fig. S1) (Table S1). For convenience, to evaluate the monitoring methods, we collected water samples from the lakeshore and thereby avoided contamination via floating gears^14^. Initially, for the individual samples, we collected 1 L of surface water in a bottle. To avoid DNA contamination, we sterilized the bottles and all other equipment used in sampling, including filtering apparatus, using 10% commercial bleach (ca. 0.6% hypochlorous acid) followed by washing with DNA-free distilled water. For the Mix_cool and Mix_freeze samples, we collected two 150-mL samples of surface water at each of the six sampling sites, which were mixed in two sterilized bottles to give two composite water samples with final volumes of 900 mL. The “field blank” sample contained 1 L of DNA-free water, which we brought to the field and treated identically to the other water samples, with the exception that it was not exposed to the external environment at the field sites. The individual, Mix_cool, and field blank water samples were stored on-site in a cooler with ice packs and transported to the laboratory in a 4°C refrigerator within 2 days. The Mix_freeze samples were stored on-site in a cooler with ice packs, transferred to a −18°C freezer within 12 h, and transported frozen at −18°C within 2 days.

### Water filtering and eDNA extraction from filter samples

Bottled water samples were vacuum-filtered through 47-mm GF/F glass fiber filters (nominal pore size: 0.7 μm; GE Healthcare, Little Chalfont, UK) in the laboratory. Following filtration, all filters were stored at −20°C prior to eDNA extraction. The field blank samples were processed in a similar manner. As an equipment control, we used 1 L of Milli-Q water to monitor contamination during the filtering of the samples from each site and during the subsequent DNA extraction.

eDNA was extracted from filters using the method developed by Uchii et al.^46^. We incubated filters by submerging in a mixed buffer comprising 400 μL of Buffer AL (Qiagen, Hilden, Germany) and 40 μL of Proteinase K (Qiagen, Hilden, Germany) using a Salivette tube (Sarstedt, Nümbrecht, Germany) at 56°C for 30 min. The Salivette tube containing filters was centrifuged at 5000 × *g* for 5 min, after which we added 220 μL of TE buffer (10 mM Tris-HCl and 1 mM EDTA, pH = 8.0) onto the filter, which was then centrifuged at 5000 × *g* for 5 min. The DNA in the eluted solution was purified using a DNeasy Blood & Tissue kit (Qiagen, Hilden, Germany), which extracted the DNA in 200 μL of Buffer AE. Samples were stored at 20°C until used for the 1st PCR assay.

### Library preparation and MiSeq sequencing

Details of the two-step PCR procedure used for Illumina MiSeq sequencing have been described previously by Fujii et al.^16^. We performed the 1st PCR using MiFish-U-F and MiFish-U-R primers^22^, which were designed to contain Illumina sequencing primer regions and 6-mer random bases, as follows:

Forward: *5’-ACACTCTTTCCCTACACGACGCTCTTCCGATCT* NNNNNN GTCGGTAAAACTCGTGCCAGC-3’;
Reverse: 5’-*GTGACTGGAGTTCAGACGTGTGCTCTTCCGATCT* NNNNNN CATAGTGGGGTATCTAATCCCAGTTTG-3’,

in which the italicized and non-italicized base sequences represent the MiSeq sequencing primers and MiFish primers, respectively, and the six random bases (N) were used to enhance cluster separation on flow cells during the initial base call calibrations on MiSeq.

We performed the 1st PCR using a 12-μL reaction volume containing 1 × PCR Buffer for KOD FX Neo polymerase (Toyobo, Osaka, Japan), 0.4 mM dNTP mix, 0.24 U KOD FX Neo polymerase, 0.3 μM of each primer, and 2 μL of template DNA. The thermocycling conditions for this step were as follows: initial denaturation at 94°C for 2 min; followed by 35 cycles of denaturation at 98°C for 10 s, annealing at 65°C for 30 s, and extension at 68°C for 30 s; and a final extension at 68°C for 5 min. The 1st PCRs were performed using eight replicates^25^, and the products of individual 1st PCR replicates were pooled and purified using AMPure XP (Beckman Coulter, Brea CA, USA) as templates for the 2nd PCR. The Illumina sequencing adapters and 8-bp identifier indices (the X sequence in the following primers) were added to the subsequent PCR process using the following forward and reverse fusion primers:

Forward: 5’-*AATGATACGGCGACCACCGAGATCTACA* XXXXXXXX ACACTCTTTCCCTACACGACGCTCTTCCGATCT-3’;
Reverse: *5’-CAAGCAGAAGACGGCATACGAGAT* XXXXXXXX GTGACTGGAGTTCAGACGTGTGCTCTTCCGATCT-3’,

in which the italicized and non-italicized base sequences represent MiSeq P5/P7 adapter and sequencing primers, respectively. The eight X bases represent dual-index sequences inserted to identify different samples^47^. We performed the 2nd PCR with 12 cycles using a 12-μL reaction volume containing 1 × KAPA HiFi HotStart ReadyMix, 0.3 μM of each primer, and 1.0 μL of the 1st PCR product. The thermocycling conditions after an initial 3-min denaturation at 95°C were as follows: denaturation at 98°C for 20 s, followed by combined annealing and extension at 72°C (shuttle PCR) for 15 s; and a final extension at the same temperature for 5 min. The 2nd PCR products were pooled in equal volumes and purified using AMPure XP.

The purified PCR products were loaded onto a 2% E-Gel SizeSelect agarose gel (Thermo Fisher Scientific, Waltham, MA, USA), and those of the target size (approximately 370 bp) were collected. The collected samples were quantified using a Qubit dsDNA HS assay kit and a Qubit 2.0 Fluorometer (Thermo Fisher Scientific). The amplicon libraries were sequenced using 2 × 250 bp paired-end sequencing on the MiSeq platform using an MiSeq v2 Reagent Kit. Note that sequencing runs were performed for a total of 415 libraries, comprising 178 libraries constructed in the present study (144 samples, 18 field and filter blanks, and 16 PCR negative controls) and 237 libraries constructed in previous research projects. MiSeq sequencing was conducted at the Department of Environmental Solution Technology, Faculty of Science and Technology, Ryukoku University. All sequence data generated have been deposited in the DNA Data Bank of Japan [DRA, Accession number: (submitted)].

### Bioinformatic analysis for MiSeq sequencing

The detailed procedures used for bioinformatics analyses have been described previously by Fujii et al.^16^. Initially, low-quality tails were trimmed from each read, and paired-end reads were then assembled. For the 5,225,947 reads thus obtained, primer sequences were removed, and identical sequences (i.e., 100% sequence similarity) were merged using UCLUST (usearch 7.0.1001)^48^. Sequences with ten or more identical reads were subjected to downstream processing. For taxonomic annotation, we conducted a local BLASTN search using BLAST 2.2.29 based on a previously established reference database of fish species for processed reads^22^. For each assessed sequence, the top BLAST hit with a sequence identity ≥98.6% was used for species detection. Note that a majority of the species were identified with a match of ≥99%. Based on the BLAST results, we identified the species and genus using previously described methods^16,20^. The sequence reads in the pipeline processes are listed in Table S2.

### Fish fauna record data

We used fish fauna data from Japanese lakes published previously by Matsuzaki et al.^7,49^. These studies assembled information on the distribution (i.e., presence or absence) of strictly freshwater fish that are intolerant of saltwater and inhabit freshwater environments for their entire life cycle, based on extensive surveys of scientific papers, monographs and books, online databases (National Survey on the Natural Environment by the Ministry of the Environment and National Censuses on River Environments by the Ministry of Land, Infrastructure, Transport and Tourism), museum specimen databases collected by the National Museum of Nature and Science, local museum specimens, prefectural and municipal reports, and gray literature, including reports of non-governmental organizations (NGOs) and universities. In the present study, using the same sources, we further added the distributions of secondary freshwater fish, which have a certain degree of salt tolerance and are occasionally able to cross narrow sea barriers, and peripheral freshwater fish, which are derived from marine ancestors and include diadromous fish that migrate between fresh and marine environments. Having compiled the distribution data, we generated a comprehensive dataset of fish fauna consisting of native and exotic fishes after extirpations and introductions. Finally, we obtained records for a total of 242 fish taxa (Table S3). To evaluate the performance of our eDNA metabarcoding survey, we assumed that the records obtained provided a reliable indication of the fish communities inhabiting the selected lakes.

We acknowledge the potential limitations associated with analyzing such compiled data, given that these data were not systematically collected. Moreover, fish were collected using a variety of apparatus and techniques (such as electrofishing, bag seines, trap nets, minnow traps, gill nets, hand nets, and fyke nets), by different entities, and within different studies with differing objectives. However, the utilization of integrated multiple sources can serve to minimize bias and summarize fish fauna in terms of temporal and spatial replication^7,48^.

### Fish trait data

We obtained details on the ecological and biological traits of the detected and recorded fish species from FishBase^50^, which was searched using species names, and obtained the related data using the “species” and “ecology” functions of the “rfishbase” package (ver. 3.1.0) on March 4, 2020. We were able to obtain sufficient data on body length (cm, including both total and standard lengths, but mainly standard length), recorded longevity in the wild (years), lateral body shape types, habitat types, and saltwater tolerance, but were unable to obtain sufficient data (for less than 12% of taxa) for certain traits, such as trophic position, and body weight, for further analysis.

### Lake morphology and characteristic data

We obtained data on lake morphology [surface area (km^2^), maximum water depth (m), mean water depth (m), and volume (km^3^)], and lake types [trophic state and water types (brackish or freshwater)] from Tanaka^51^ (Table S1).

### Statistical analyses

To evaluate the performance of eDNA metabarcoding with respect to detecting species, we used “percentage record covered (%)” as a measure of the proportion of eDNA-detected species matching species recorded in the target lakes, which was calculated from the number of eDNA-detected species matched to the fish records per total number of recorded species in the dataset.

All statistical analyses and graphic preparations were performed using R ver. 3.6.0^52^, and statistical values were evaluated at a significance level of α = 0.05.

To compare eDNA metabarcoding and species record data, we compared taxonomic levels in the species list compiled based on visual surveys with those in the lists compiled using eDNA metabarcoding data (Table S1, S2) with reference to previous studies that have used MiFish primers^16,20^. To confirm that the sequencing depth was sufficient to detect fish diversity in the samples, we used the “rarefy” and “rarecurve” functions of the “vegan” package (ver. 2.5.6) (Fig. S2). Thus, we used raw data for further analyses without rarefying the data.

We examined differences in the number of detected fish taxa and percentage records covered evaluated by eDNA metabarcoding among the sampling methods (individual, Mix_cool, Mix_freeze, and field and negative controls) using nested analysis of variance (nested ANOVA) based on LMMs with “lake” as the nested (random) factor, using the “anova.lme” and “lme” functions of the “nlme” package (ver. 3.1.139), respectively. Tukey’s post hoc test was used for multiple comparisons for least-squares means using the “lsmeans” function of the “lsmeans” package (ver. 2.30.0).

Differences in fish traits, body length (numeric data), habitats, lateral body shape types, and saltwater tolerance (categorical data) between high frequency of species detection (eDNA metabarcoding vs. species record) among the sampling methods (individual vs. Mix_cool vs. Mix_freeze) were examined using nested ANOVA and an LMM as described above. Prior to the GLM analyses, we calculated a variance inflation factor (VIF) to check the collinearity of the explanatory factors of the GLMs. We found that the maximum VIF value was 1.23 and that all VIF values were less than 5.

We also performed GLM analysis to evaluate the relationships between the percentage record covered of the species record and the explanatory factors of lake ecosystems, including lake latitude and altitude, morphology (surface area, maximum water depth, mean water depth, and volume), trophic state (eutrophic vs. meso- and oligotrophic), and water types (freshwater vs. brackish) using the “glm” function. We set “binomial” as the error distribution of the GLMs and added the total number of species records as the offset factor. In this case, we found that the maximum VIF value was 49.7, indicating that collinearity among the factors could potentially influence the parameter estimations in the GLMs. After removing the explanatory factors with a VIF value >5 to reduce the collinearity effect on the GLMs, we finally performed the GLMs using latitude, surface area, mean water depth, trophic state, and water types for all sampling methods.

We performed indicator taxa analysis^53^ to determine those taxa showing significantly different frequencies between the eDNA metabarcoding and species record species lists. The analysis was performed using the “signassoc” function of the “indicspecies” package (ver. 1.7.6) based on presence/absence data^53^. We used mode = 1 (group-based) and calculated the P values with 999 permutations after applying Sidak’s correction for multiple testing.

Differences in community compositions were visualized using non-metric multidimensional scaling (NMDS) with 500 separate runs of real data. The community dissimilarity for NMDS was calculated using incidence-based Jaccard indices. We also calculated NMDS stress to confirm the representation of the NDMS ordination and evaluated differences in community structures between sampling methods and sites using PERMANOVA, for which we used an incidence-based Jaccard similarity matrix and calculated the statistical values with 999 permutations. For NMDS and PERMANOVA analyses, we used the “metaMDS” and “Adonis” functions of the “vegan” package (ver. 2.5.6), respectively.

## Acknowledgments

This study was supported by the Environment Research and Technology Development Fund (4-1602, 4-1705, 4-2004) of the Environmental Restoration and Conservation Agency, Japan, and JST-CREST (JPMJCR13A2).

## Data availability

All data obtained from MiSeq sequencing are available in DRA [Accession number: DRA (submitted)], and all data used, including all species detected based on MiSeq sequencing, the recorded species data, and lake data used for analysis, are shown in the Supplemental tables.

## Author contributions

HD designed the study, and HA, KI, AK, KK, NM, T Mitsuzuka, TT, KT, NU, TW, and KY contributed to field sampling. SSM contributed to preparing the dataset of the fish presence records. S. Matsuoka, MN, HD, HS, HY, S Matsuhashi, SY, T Minamoto, and MM contributed to laboratory and molecular experiments. HD, S Matsuoka, and SSM analyzed the data. HD, S Matsuoka, SSM, and MM wrote the initial draft of the manuscript. All other authors critically reviewed the manuscript.

## Notes

### Competing Interest Statement

The authors have declared no competing interest.

## References

1. Dudgeon, D. et al. Freshwater biodiversity: importance, threats, status and conservation challenges. Biol. Rev. 81, 163–182 (2006).

2. Flitcroft, R., Cooperman, M. S., Harrison, I. J., Juffe-Bignoli, D. & Boon, P. J. Theory and practice to conserve freshwater biodiversity in the Anthropocene. Aquat. Cons. Mar. Freshw. Ecosyst. 29, 1013–1021 (2019).

3. Strayer, D. L. & Dudgeon, D. Freshwater biodiversity conservation: recent progress and future challenges. J. North Am. Benthol. Soc. 29, 344–358 (2010).

4. Collen, B. et al. Global patterns of freshwater species diversity, threat and endemism. Glob. Ecol. Biog. 23, 40–51 (2014).

5. Gardner, J. L., Amano, T., Sutherland, W. J., Joseph, L. & Peters, A. Are natural history collections coming to an end as time-series? Front. Ecol. Env. 12, 436–438 (2014).

6. Schindler, D. E. & Hilborn, R. Prediction, precaution, and policy under global change. Science 347, 953–954 (2015).

7. Matsuzaki, S. S., Sasaki, T. & Akasaka, M. Invasion of exotic piscivores causes losses of functional diversity and functionally unique species in Japanese lakes. Freshw. Biol. 61, 1128–1142 (2016a).

8. Socolar, J. B., Gilroy, J. J., Kunin, W. E. & Edwards, D. P. How should beta-diversity inform biodiversity conservation? Trends Ecol. Evol. 31, 67–80 (2016).

9. Nishihiro, J. et al. Heterogeneous distribution of a floating-leaved plant, Trapa japonica, in Lake Mikata, Japan, is determined by limitations on seed dispersal and harmful salinity levels. Ecol. Res. 29, 981–989 (2014).

10. Takahara, T., Minamoto, T., Yamanaka, H., Doi, H. & Kawabata, Z. Estimation of fish biomass using environmental DNA. PloS ONE 7, e35868 (2012).

11. Rees, H. C., Maddison, B. C., Middleditch, D. J., Patmore, J. R. & Gough, K. C. Review: the detection of aquatic animal species using environmental DNA–a review of eDNA as a survey tool in ecology. J. Appl. Ecol. 51, 1450–1459 (2014).

12. Goldberg, C. S., Strickler, K. M. & Pilliod, D. S. Moving environmental DNA methods from concept to practice for monitoring aquatic macroorganisms. Biol. Cons. 183, 1–3 (2015).

13. Thomsen, P. F. & Willerslev, E. Environmental DNA–an emerging tool in conservation for monitoring past and present biodiversity. Biol. Cons. 183, 4–18 (2015).

14. Doi, H. et al. Environmental DNA analysis for estimating the abundance and biomass of stream fish. Freshw. Biol. 6, 30–39 (2017a).

15. Yamamoto, S. et al. Environmental DNA metabarcoding reveals local fish communities in a species-rich coastal sea. Sci. Rep. 7, 1–12 (2017).

16. Fujii, K., Doi, H., Matsuoka, S., Nagano, S., Sato, H. & Yamanaka, H. Environmental DNA metabarcoding for fish community analysis in backwater lakes: A comparison of capture methods. PLoS ONE 14, e0210357 (2019).

17. Bylemans, J. et al. Monitoring riverine fish communities through eDNA metabarcoding: determining optimal sampling strategies along an altitudinal and biodiversity gradient. Metabar. Metagen. 2, e30457 (2018).

18. Deiner, K., Fronhofer, E. A., Mächler, E., Walser, J. C. & Altermatt, F. Environmental DNA reveals that rivers are conveyer belts of biodiversity information. Nat. Comm. 7, 12544 (2016).

19. Deiner, K. et al. Environmental DNA metabarcoding: Transforming how we survey animal and plant communities. Mol. Ecol. 26, 5872–5895 (2017).

20. Sato, H., Sogo, Y., Doi, H. & Yamanaka, H. Usefulness and limitations of sample pooling for environmental DNA metabarcoding of freshwater fish communities. Sci. Rep. 7, 14860 (2017).

21. Taberlet, P., Coissac, E., Pompanon, F., Brochmann, C. & Willerslev, E. Towards next-generation biodiversity assessment using DNA metabarcoding. Mol. Ecol. 21, 2045–2050 (2012).

22. Miya, M. et al. MiFish, a set of universal PCR primers for metabarcoding environmental DNA from fishes: detection of more than 230 subtropical marine species. Royal Soc. Open Sci. i, 150088 (2015).

23. Nakagawa, H., Yamamoto, S., Sato, Y., Sado, T., Minamoto, T. & Miya, M. Comparing local- and regional-scale estimations of the diversity of stream fish using eDNA metabarcoding and conventional observation methods. Freshw. Biol. 63, 569–580 (2018).

24. Collins, M. K. et al. Searching for a Salamander: Distribution and Habitat of the Mudpuppy (*Necturus maculosus*) in Southeast Ohio Using eDNA as a Rapid Assessment Technique. Ame. Midland Nat, 182, 191–202 (2019)..

25. Doi, H., Fukaya, K., Oka, S. I., Sato, K., Kondoh, M. & Miya, M. Evaluation of detection probabilities at the water-filtering and initial PCR steps in environmental DNA metabarcoding using a multispecies site occupancy model. Sci. Rep. 9, 3581 (2019).

26. Cilleros, K. et al. Unlocking biodiversity and conservation studies in high-diversity environments using environmental DNA (eDNA): A test with Guianese freshwater fishes. Mol. Ecol. Res. 19, 27–46 (2019).

27. Li, J. et al. Ground-truthing of a fish-based environmental DNA metabarcoding method for assessing the quality of lakes. J. Appl. Ecol. 56, 1232–1244 (2019).

28. Jerde, C. L., Wilson, E. A. & Dressler, T. L. Measuring global fish species richness with eDNA metabarcoding. Mol. Ecol. Res. 19, 19–22 (2019).

29. Stat, M. et al. Ecosystem biomonitoring with eDNA: metabarcoding across the tree of life in a tropical marine environment. Sci. Rep. 7, 1–11 (2017).

30. Maruyama, A. et al. The release rate of environmental DNA from juvenile and adult fish. PLoS One, 9, e114639 (2014).

31. Pont, D. et al. The future of fish-based ecological assessment of European rivers: from traditional EU Water Framework Directive compliant methods to eDNA metabarcoding-based approaches. J. Fish Biol. 1–13 (2019).

32. Deiner, K., Walser, J. C., Mächler, E. & Altermatt, F. Choice of capture and extraction methods affect detection of freshwater biodiversity from environmental DNA. Biol. Cons. 183, 53–63 (2015).

33. Takahara, T., Minamoto, T. & Doi, H. Using environmental DNA to estimate the distribution of an invasive fish species in ponds. PLoS ONE 8, e56584 (2013).

34. Nilsson, A. N., Elmberg, J. & Sjoberg, K. Abundance and species richness patterns of predaceous diving beetles (Coleoptera, Dytiscidae) in Swedish lakes. J. Biogeogr. 82, 197–206 (1994).

35. Bista, I. et al. Annual time-series analysis of aqueous eDNA reveals ecologically relevant dynamics of lake ecosystem biodiversity. Nat. Comm. 8, 14087 (2017).

36. Doi, H. et al. Water sampling for environmental DNA surveys by using an unmanned aerial vehicle. Limnol. Ocean. Meth. 15, 939–944 (2017b).

37. Dodson, S. I., Arnott, S. E. & Cottingham, K. L. The relationship in lake communities between primary productivity and species richness. Ecology 81, 2662–2679 (2000).

38. Post, D. M., Pace, M. L. & Hairston, N. G. Ecosystem size determines food-chain length in lakes. Nature 405, 1047–1049 (2000).

39. Wetzel R. G. Limnology: lake and river ecosystems. 3rd edition. Gulf professional publishing, Huston, USA (2001)

40. Harper, L. R. et al. Prospects and challenges of environmental DNA (eDNA) monitoring in freshwater ponds. Hydrobiologia 826, 25–41 (2019).

41. Guillera-Arroita, G., Lahoz-Monfort, J. J., van Rooyen, A. R., Weeks, A. R. & Tingley, R. Dealing with false-positive and false-negative errors about species occurrence at multiple levels. Meth. Ecol. Env. 8, 1081–1091 (2017).

42. Lahoz-Monfort, J. J., Guillera-Arroita, G. & Tingley, R. Statistical approaches to account for false - positive errors in environmental DNA samples. Mol. Ecol. Res. 16, 673–685 (2016).

43. Connor, E. F. & McCoy, E. D. The statistics and biology of the species-area relationship. Am. Nat. 113, 791–833 (1979).

44. Ugland, K. I., Gray, J. S. & Ellingsen, K. E. The species–accumulation curve and estimation of species richness. J. Anim. Ecol. 72, 888–897 (2003).

45. Sigsgaard, E. E. et al. Using vertebrate environmental DNA from seawater in biomonitoring of marine habitats. Cons. Biol. 34, 697–710 (2020).

46. Uchii, K., Doi, H. & Minamoto, T. A novel environmental DNA approach to quantify the cryptic invasion of non-native genotypes. Mol. Ecol. Res. 16, 415–422 (2016).

47. Hamady, M., Walker, J. J., Harris, J. K., Gold, N. J. & Knight, R. Error-correcting barcoded primers for pyrosequencing hundreds of samples in multiplex. Nat. Meth. 5, 235–237 (2008).

48. Edgar, R. C. Search and clustering orders of magnitude faster than BLAST. Bioinformatics 26, 2460–2461 (2010).

49. Matsuzaki, S.S. et al. Biodiversity of freshwater fish and aquatic macrophytes in Japanese lakes: a broad assessment. Jap. J. Cons. Ecol. 21, 155–165 (2016b). (in Japanese)

50. Froese, R. & Pauly D. (Eds.). (2019, December Day). FishBase [Database]. Retrieved from www.fishbase.org

51. Tanaka M. The Lakes in Japan, 530 pp. in Japanese, Nagoya University Press, Nagoya, (1992).

52. R Core Team. R: A language and environment for statistical computing. R Foundation for Statistical Computing, Vienna, Austria (2019). Retrieved from https://www.R-project.org/

53. Cáceres, M. D. & Legendre, P. Associations between species and groups of sites: indices and statistical inference. Ecology 90, 3566–3574 (2009).

