## Supplemental Materials for "Effects of species traits and ecosystem characteristics on species detection by eDNA metabarcoding in lake fish communities"

Table S1 Sampling date, lake morphology, types and location of the 17 study lakes. CSV file (LakeeDNA-SEM-TableS1.csv).

Table S2 The sequence read of the deta processing and taxon assignment by MiFish pipeline. CSV file (LakeeDNA-SEM-TableS2.csv).

Table S3 Fish taxa list recorded by species record. CSV file (LakeeDNA-SEM-TableS3.csv).

Table S4 Detected fish taxa list evaluated by eDNA metabarcoding (eDNA). CSV file (LakeeDNA-SEM-TableS4.csv). Note: we excluded *Pagrus major* from the statistical analysis.

Table S5 Nested ANOVA with LMM and Tukey post-hoc test on least-squares means for number of detected taxa, the percentage covered of record evaluated by eDNA metabarcoding in the three sampling methods.

Table S6 Nested ANOVA with LMM for the differences in fish traits, body length (numeric), habitats, body shape types, and saltwater tolerance (categories) between highly frequency of species-detection (eDNA metabarcoding vs species record) with the among the sampling methods (Individual vs Mix_cool vs Mix_freeze).

Table S7 The GLM results for the percentage covered of species records with the explanatory factors, including latitude, surface area, mean water depth, trophic state (eutrophic vs meso- and oligotrophic) and water types (brackish or freshwater), in three sampling methods.

Table S8 Indicator taxa analysis to determine which taxa had significantly different frequency between the both communities evaluated by eDNA metabarcoding and species records. Best means the frequently detected methods. The P values was calculated with 999 permutations after Sidak's correction for multiple testing. CSV file (LakeeDNA-SEM-TableS8.csv).

Table S9 PERMANOVA results for community structures evaluated by eDNA metabarcoding and historical record.

Figure S1 Map for the sampling sites in each study lake. (Fig. S1.pdf).

Figure S2 The relationship between number of sequence reads and the number of detected taxa, i.e., rarefaction curves.

Figure S3 Number of detected fish taxa by eDNA metabarcoding with different sampling methods. "Total" means total number of detected fish taxa by eDNA metabarcoding.

Figure S4 The box and violin plot for the detected fish taxa by eDNA metabarcoding among the sampling methods and controls. Blank and NC means filed blank (control) and PCR negative controls, respectively. Boxes and bar in the box plot indicate median ± inter-quartiles, and ±1.5 × inter-quartiles. The smooth lines indicate the distribution of the data using a violin plot. The violin plot outlines illustrate kernel probability density, i.e. the width of the shaded area represents the proportion of the data located.

Figure S5 The box plot for the detected fish taxa by eDNA metabarcoding in the lakes (red). Blue and green points indicate Mix_cool and Mix_freeze data, respectively. Boxes and bar in the box plot indicate median ± inter-quartiles, and ±1.5 × inter-quartiles.

Figure S6 Percentage of record covered by eDNA metabarcoding (%) on species record in each lake.

Figure S7 The relationships between the percentage of record covered by eDNA metabarcoding (%) on species dataset with the factors without the final GLMs for the sampling methods (red=Individual, blue=Mix_cool, and green=Mix_freeze). The solid and gray area indicate the regression line from the GLM result and the 95% CI, respectively.

Figure S8 NMDS ordination plot for fish communities of species record and eDNA metabarcoding using Individual samples. NMDS stress was 0.206.

Figure S9 NMDS ordination plot for fish communities of species record and eDNA metabarcoding using Mix_freeze samples. NMDS stress was 0.179.

Table S5 Nested ANOVA with LMM and Tukey post-hoc test on least-squares means for number of detected taxa, percentage covered record evaluated by eDNA metabarcoding in the three sampling methods.

a) Nested ANOVA with LMM for number of detected taxa with blank samples

| Parameters | Df | residual Df | F | P |
|---|---|---|---|---|
| Sampling methods: lake | 4 | 132 | 24.61 | <0.0001 |
| (Intercept) | 1 | 132 | 101.11 | <0.0001 |

b) Nested ANOVA with LMM for number of detected taxa

| Parameters | Df | residual Df | F | P |
|---|---|---|---|---|
| Sampling methods: lake | 2 | 120 | 5.22 | 0.0066 |
| (Intercept) | 1 | 120 | 108.03 | <0.0001 |

c) Tukey test results for number of detected taxa in different sampling methods. CI means confidential interval.

| Pairs | Estimate | SE | Df | t ratio | P |
|---|---|---|---|---|---|
| Mix_cold-Individual | -4.3 | 1.52 | 120 | -2.83 | 0.0151 |
| Mix_freeze-Individual | -2.9 | 1.48 | 120 | -1.96 | 0.1274 |
| Mix_freeze-Mix_cold | 1.41 | 1.96 | 120 | 0.72 | 0.7545 |

Table S6 Nested ANOVA with LMM for the differences in fish traits, body length (numeric), habitats, body shape types, and saltwater tolerance (categories) between highly frequency of species-detection (eDNA metabarcoding vs species record) with the among the sampling methods (Individual vs Mix_cool vs Mix_freeze).

a) Body length

| Parameters | Df | residual Df | F | P |
|---|---|---|---|---|
| (Intercept) | 1 | 1395 | 50.2623 | <.0001 |
| eDNA vs Record | 1 | 1395 | 26.94523 | <.0001 |
| Sampling method | 2 | 1395 | 0.00955 | 0.9905 |

b) Habitat

| Parameters | Df | residual Df | F | P |
|---|---|---|---|---|
| (Intercept) | 1 | 16 | 3.1777 | 0.0936 |
| eDNA vs Record | 1 | 16 | 501.9759 | <.0001 |
| Sampling method | 2 | 16 | 0.0401 | 0.9608 |

c) Saltwater tolerance

| Parameters | Df | residual Df | F | P |
|---|---|---|---|---|
| (Intercept) | 1 | 6 | 59.00676 | 0.0003 |
| eDNA vs Record | 1 | 6 | 26.54074 | 0.0021 |
| Sampling method | 2 | 2 | 0.81355 | 0.5514 |

d) Body shape type

| Parameters | Df | residual Df | F | P |
|---|---|---|---|---|
| (Intercept) | 1 | 10 | 1.5779 | 0.2376 |
| eDNA vs Record | 1 | 10 | 37.33217 | 0.0001 |
| Sampling method | 2 | 10 | 0.06715 | 0.9355 |

Table S7 GLM results for the percentage covered of species records with the explanatory factors, including latitude, surface area, mean water depth, trophic state (eutrophic vs meso- and oligotrophic) and water types.

a) Percentage covered of species record for Individual

| | Estimate | SE | t value | P |
|---|---|---|---|---|
| (Intercept) | -149.42 | 94.21 | -1.59 | 0.144 |
| Latitude | 2.36 | 2.13 | 1.11 | 0.293 |
| $Log_{10}$ Surface area | -36.75 | 11.19 | -3.28 | 0.008 |
| $Log_{10}$ mean water depth | -2.08 | 21.27 | -0.10 | 0.924 |
| Trophic state | 48.32 | 32.87 | 1.47 | 0.172 |
| Water type | 15.89 | 16.71 | 0.95 | 0.364 |

b) Percentage covered of species record for Mix_cool

| | Estimate | SE | t value | P |
|---|---|---|---|---|
| (Intercept) | -143.13 | 90.57 | -1.58 | 0.145 |
| Latitude | 2.41 | 2.04 | 1.18 | 0.265 |
| $Log_{10}$ Surface area | -36.10 | 10.76 | -3.35 | 0.007 |
| $Log_{10}$ mean water depth | 3.48 | 20.45 | 0.17 | 0.868 |
| Trophic state | 42.80 | 31.60 | 1.35 | 0.205 |
| Water type | 13.42 | 16.07 | 0.84 | 0.423 |

c) Percentage covered of species record for Mix_freeze

| | Estimate | SE | t value | P |
|---|---|---|---|---|
| (Intercept) | -149.69 | 96.65 | -1.55 | 0.153 |
| Latitude | 2.51 | 2.18 | 1.15 | 0.277 |
| $Log_{10}$ Surface area | -32.49 | 11.48 | -2.83 | 0.018 |
| $Log_{10}$ mean water depth | 5.90 | 21.82 | 0.27 | 0.793 |
| Trophic state | 39.49 | 33.72 | 1.17 | 0.269 |
| Water type | 13.31 | 17.14 | 0.78 | 0.456 |

Table S8 PERMANOVA results for community structures evaluated by eDNA metabarcoding and species record.

a) Individual

| | Df | Sums of Sqs | Mean Sqs | F. Model | $R^2$ | P |
|---|---|---|---|---|---|---|
| Lake | 17 | 6.07 | 0.36 | 2.72 | 0.68 | < 0.0001 |
| eDNA vs record | 1 | 0.67 | 0.67 | 5.14 | 0.08 | < 0.0001 |
| Residuals | 17 | 2.23 | 0.13 | 0.25 | | |
| Total | 35 | 8.97 | 1.00 | | | |

b) Mix_cool

| | Df | Sums of Sqs | Mean Sqs | F. Model | $R^2$ | P |
|---|---|---|---|---|---|---|
| Lake | 17 | 5.32 | 0.33 | 2.21 | 0.59 | < 0.0001 |
| eDNA vs record | 1 | 1.30 | 1.30 | 8.62 | 0.14 | < 0.0001 |
| Residuals | 16 | 2.41 | 0.15 | 0.27 | | |
| Total | 33 | 9.02 | 1.00 | | | |

c) Mix_freeze

| | Df | Sums of Sqs | Mean Sqs | F. Model | $R^2$ | P |
|---|---|---|---|---|---|---|
| Lake | 17 | 5.93 | 0.35 | 2.13 | 0.60 | < 0.0001 |
| eDNA vs record | 1 | 1.24 | 1.24 | 7.57 | 0.12 | < 0.0001 |
| Residuals | 17 | 2.78 | 0.16 | 0.28 | | |
| Total | 35 | 9.96 | 1.00 | | | |

a) Individual

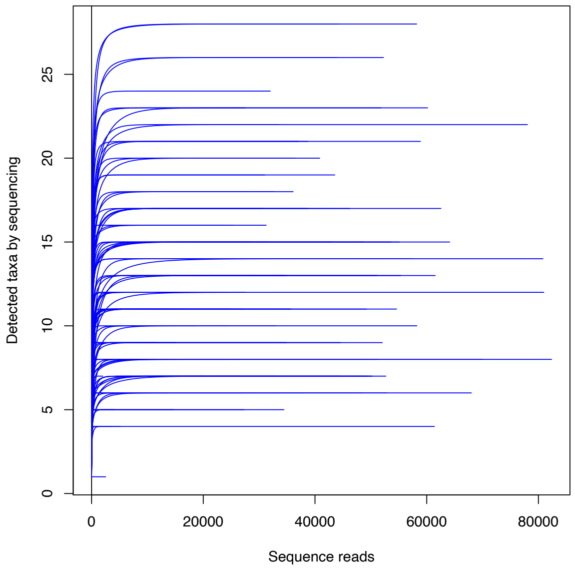


b) Mix_cool

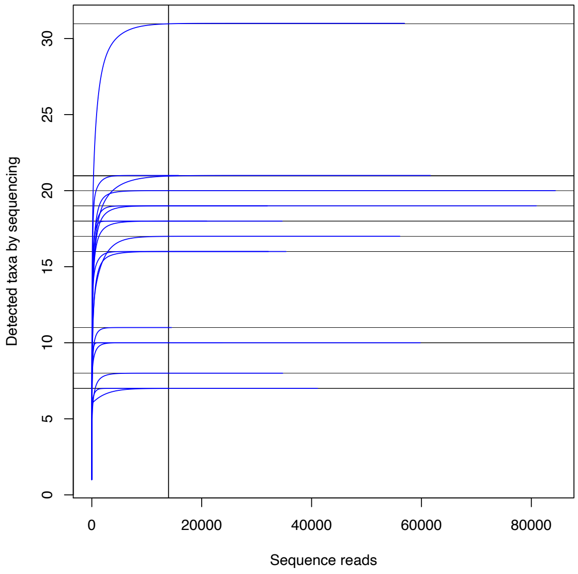

c) Mix_freeze

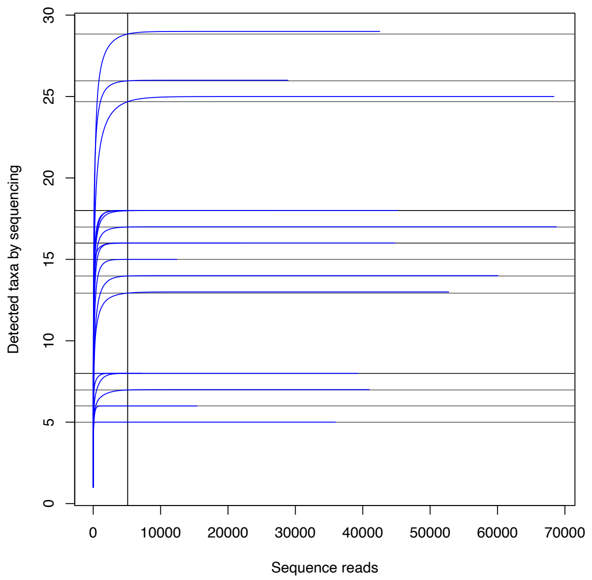


Figure S2 The relationship between number of sequence reads and the number of detected taxa, i.e., rarefaction curves.

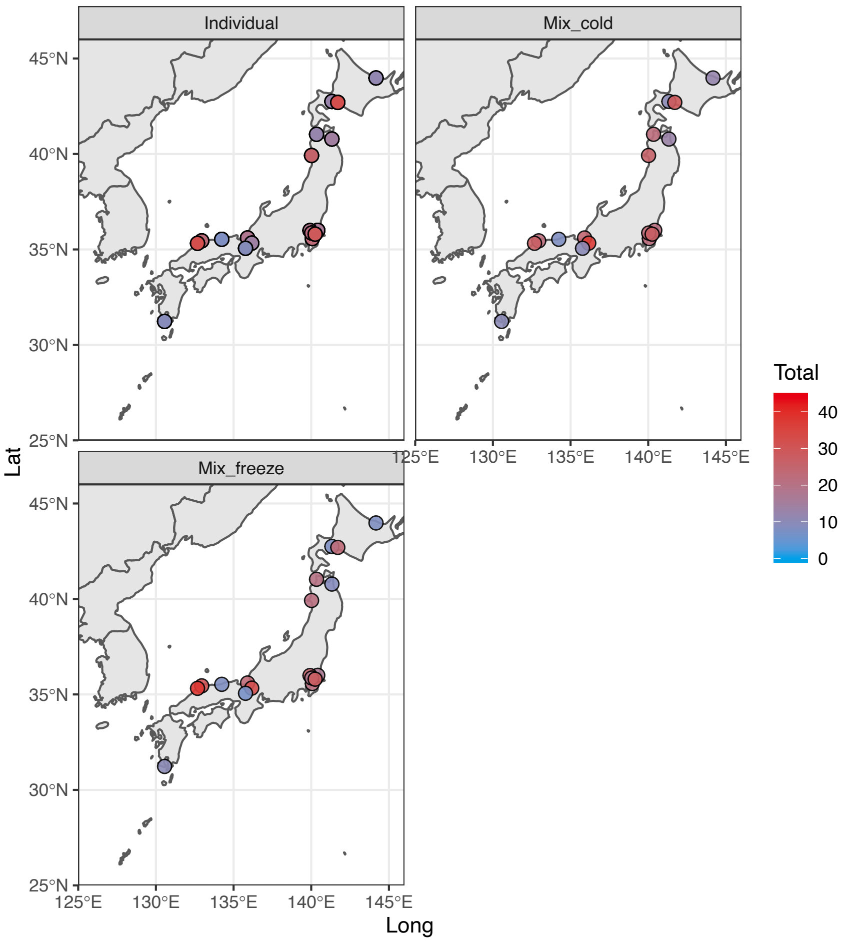


Figure S3 Number of detected fish taxa by eDNA metabarcoding with different sampling methods. "Total" means total number of detected fish taxa by eDNA metabarcoding.

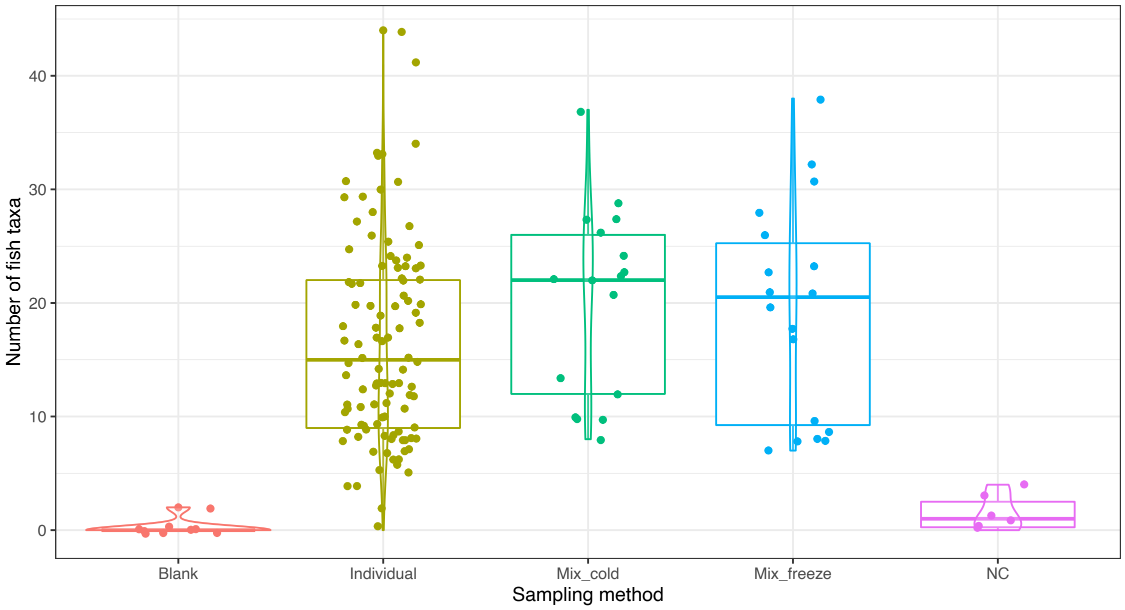


Figure S4 The box and violin plot for the detected fish taxa by eDNA metabarcoding among the sampling methods and controls. Blank and NC means filed blank (control) and PCR negative controls, respectively. Boxes and bar in the box plot indicate median ± inter-quartiles, and ±1.5 × inter-quartiles. The smooth lines indicate the distribution of the data using a violin plot. The violin plot outlines illustrate kernel probability density, i.e. the width of the shaded area represents the proportion of the data located.

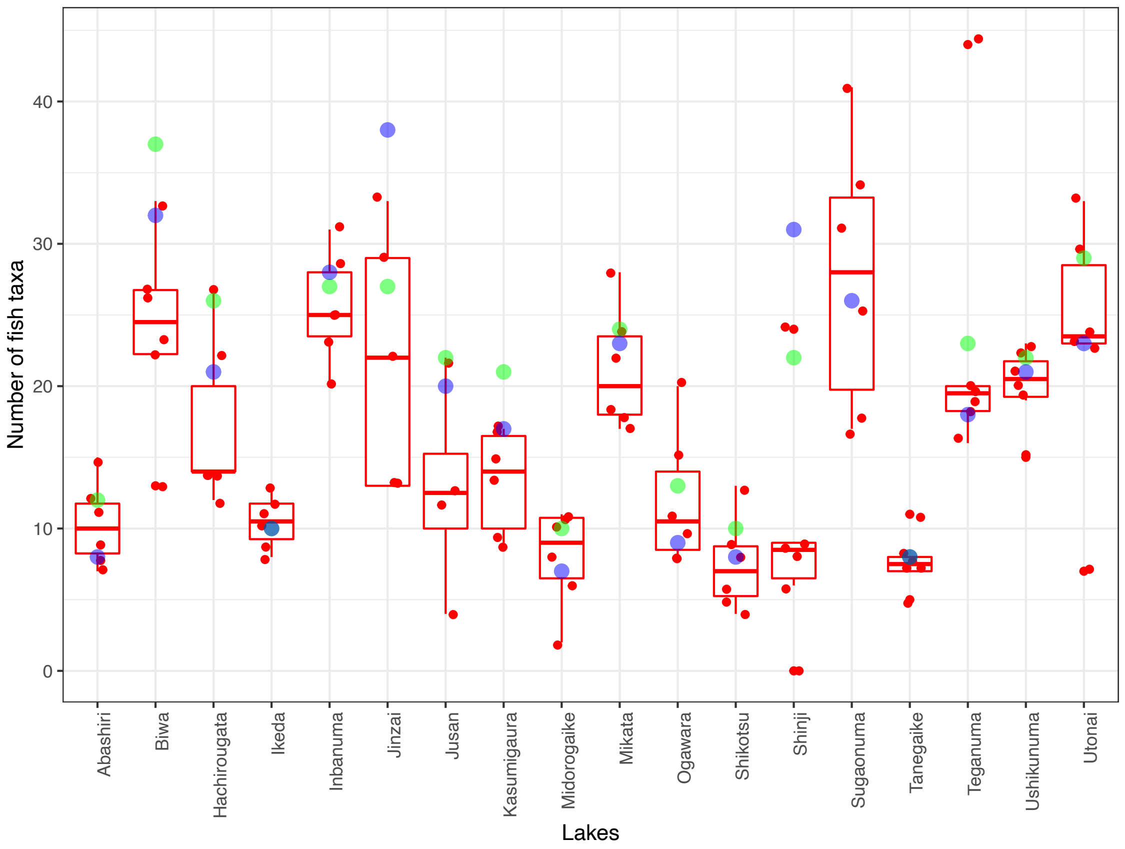


Figure S5 The box plot for the detected fish taxa by eDNA metabarcoding in the lakes (red). Blue and green points indicate Mix_cool and Mix_freeze data, respectively. Boxes and bar in the box plot indicate median ± inter-quartiles, and ±1.5 × inter-quartiles.

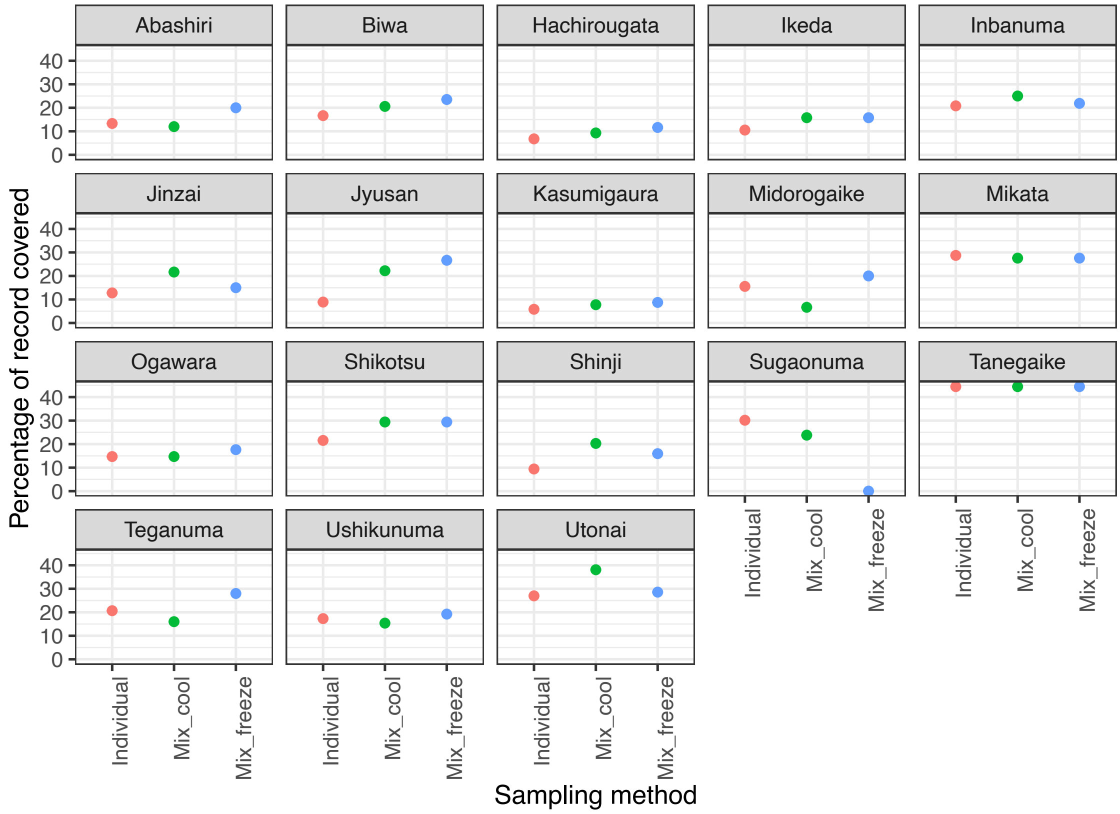


Figure S6 Percentage of record covered by eDNA metabarcoding (%) on species record in each lake.

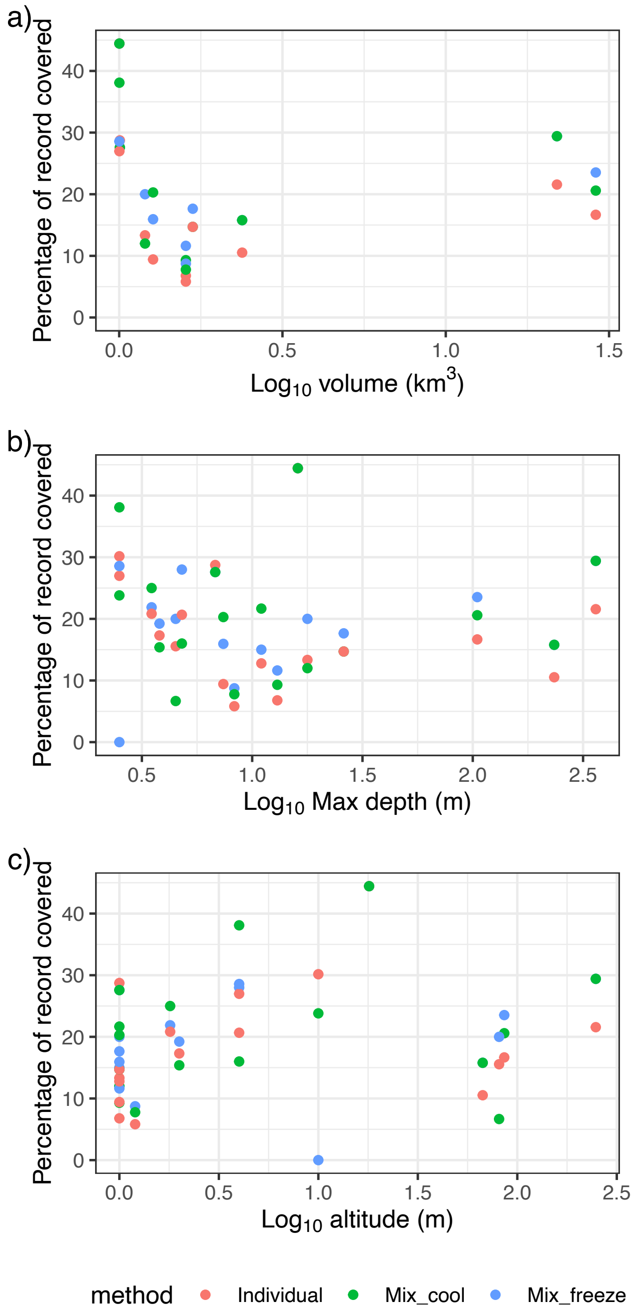


Figure S7 The relationships between the percentage of record covered by eDNA metabarcoding (%) on species dataset with the factors without the final GLMs for the sampling methods (red=Individual, blue=Mix_cool, and green=Mix_freeze). The solid and gray area indicate the regression line from the GLM result and the 95% CI, respectively.

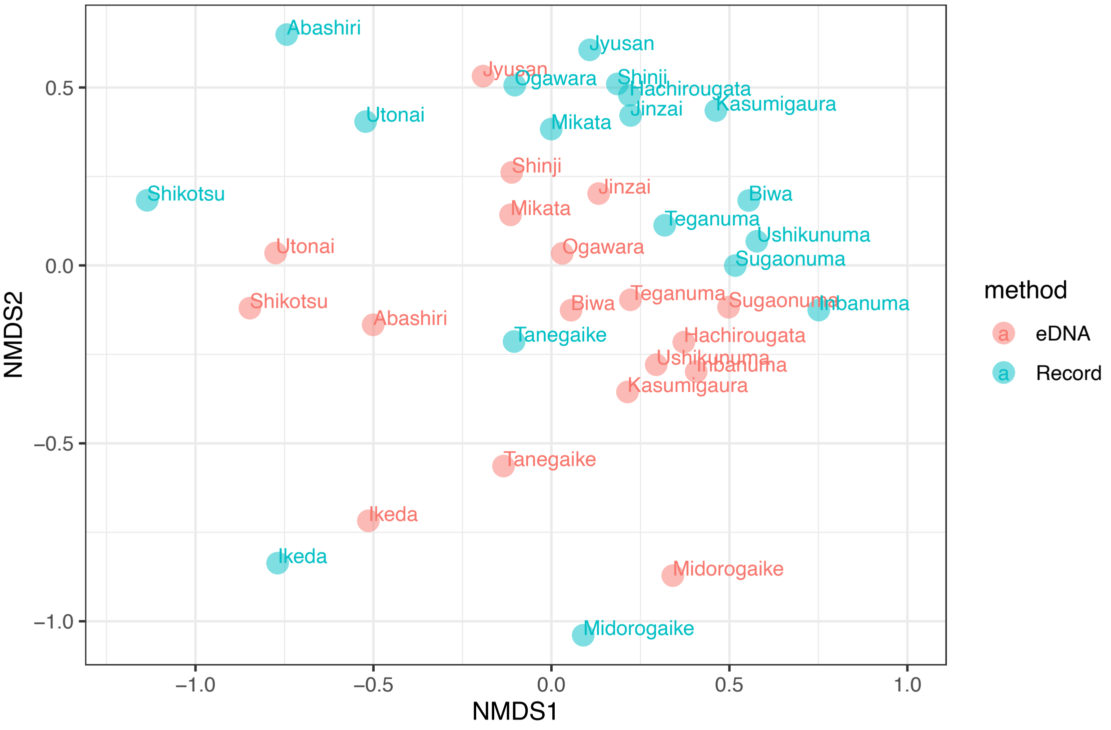


Figure S8 NMDS ordination plot for fish communities of species record and eDNA metabarcoding using Individual samples. NMDS stress was 0.206.

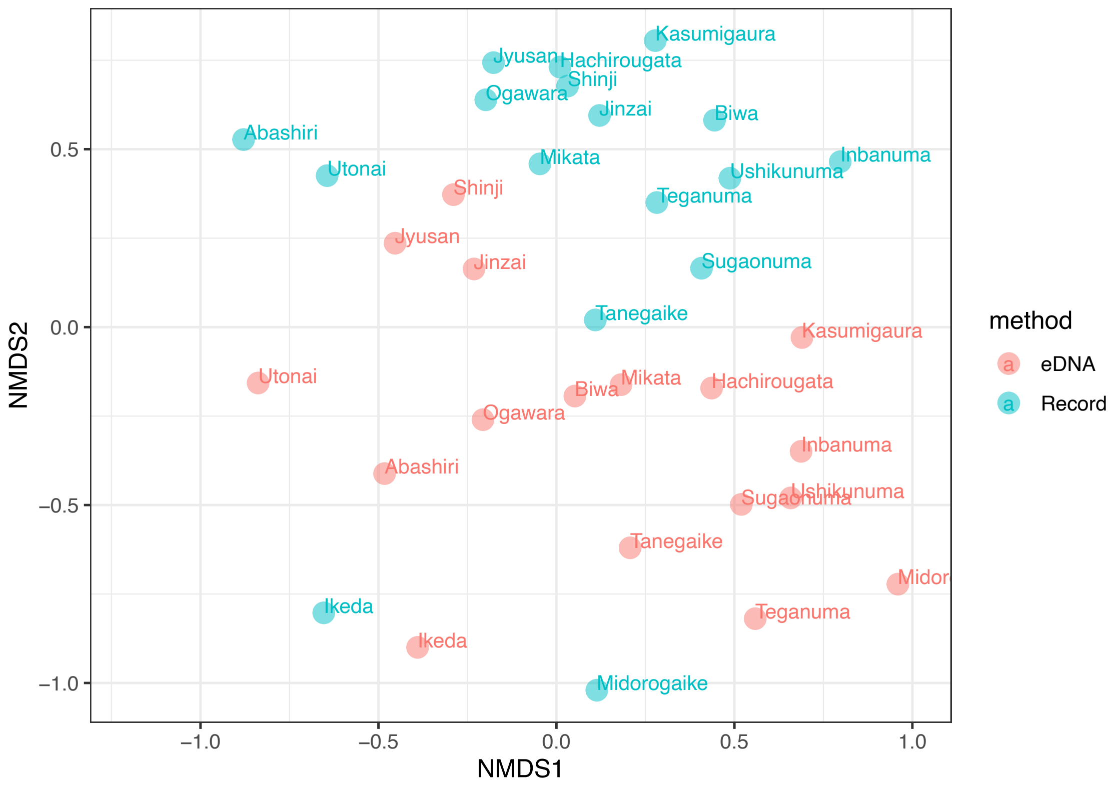


Figure S9 NMDS ordination plot for fish communities of species record and eDNA metabarcoding using Mix_freeze samples. NMDS stress was 0.179.
